# Single-cell RNA Expression of SARS-CoV-2 Cell Entry Factors in Human Endometrium during Preconception

**DOI:** 10.1101/2020.09.14.296806

**Authors:** Felipe Vilella, Wanxin Wang, Inmaculada Moreno, Stephen R. Quake, Carlos Simon

## Abstract

We investigated potential SARS-CoV-2 tropism in human endometrium by single-cell RNA-sequencing of viral entry-associated genes in healthy women. Percentages of endometrial cells expressing *ACE2, TMPRSS2, CTSB*, or *CTSL* were <2%, 12%, 80%, and 80%, respectively, with 0.7% of cells expressing all four genes. Our findings imply low efficiency of SARS-CoV-2 infection in the endometrium before embryo implantation, providing information to assess preconception risk in asymptomatic carriers.

## Introduction

The coronavirus disease 2019 (COVID-19) pandemic produced by severe acute respiratory syndrome coronavirus 2 (SARS-CoV-2) has impacted public health worldwide, including through potential risks to couples aiming to conceive or already pregnant^1^. SARS-CoV-2 preferentially infects cells in the respiratory tract, heart, liver, brain, and kidneys^2^, but knowledge of its direct tropism for non-vital reproductive organs that involve human pregnancy and neonatal heath is limited.

The endometrium is the mucosa of the human uterus. This tissue acts as the maternal interface and possible infectious gatekeeper during embryo implantation, placentation, and fetal development. Evidence suggests the existence of placental infection with SARS-CoV-2 during the second trimester of pregnancy in a woman with confirmed COVID-19^3^ as well as potential vertical transmission in ~9% of newborns from mothers infected with SARS-CoV-2^4,5^. These data implicate intrauterine passage of the virus from the uterus/endometrium to the placenta and then the fetus. However, it remains unknown whether the virus could be transmitted from the maternal endometrium to the embryo during early development, which might entail teratogenic effects.

SARS-CoV-2 uses angiotensin-converting enzyme 2 (ACE2) as the main receptor for entry into host cells^6^, suggesting cells expressing ACE2 are most susceptible to viral infection. While lung is considered the main organ site for infection, single-cell RNA-sequencing (scRNAseq) generated a “risk map” of different organs, revealing that heart, esophagus, kidney, and bladder are vulnerable to SARS-CoV-2 infection due to *ACE2* expression^7^. ACE2 is a member of the renin-angiotensin-aldosterone system (RAAS) that regulates blood pressure by direct action of angiotensin II on the cardiovascular system as well as by aldosterone on water-electrolyte balance^8^. However, ACE2 expression alone is insufficient for cell entry of SARS-CoV-2. After viral-host tropism and adhesion of SARS-CoV-2 S protein to ACE2 on the cell surface, priming of S protein between S1 and S2 units is essential for fusion to the cell membrane and viral entry into the cell. This cleavage is efficiently performed by host serine protease TMPRSS2, but in the absence of this protein, cathepsins (CTS) B and L can also cleave S protein to facilitate viral entry. The essential role of these proteases in SARS-CoV-2 internalization has been confirmed by infection blockade after chemical treatment of lung cells with camostat mesylate or E-64d, which inhibit TMPRSS2 or CTSB/L, respectively^6^.

Human endometrium expresses angiotensins 1–7, angiotensin receptor MAS, and ACE2. *ACE2* mRNA is expressed more abundantly in epithelial than stromal cells and in secretory versus proliferative phases^9^. During pregnancy, the human decidua shows abundant expression of RAAS proteins—prorenin (REN), prorenin receptor (ATP6AP2), AGT, ACE1, ACE2, AGTR1, and MAS—as demonstrated by qPCR in specimens collected at term either before labor (after cesarean section delivery) or after spontaneous labor^10^. Moreover, RAAS components including ACE2 are detected in decidua and fetal membranes in human placenta, with speculative roles in trophoblast invasion and angiogenesis^11,12^.

## Results

Clear evidence for the existence of SARS-CoV-2 cell entry machinery in the human endometrium during preconception is important to understand the risk of viral infection in this key reproductive organ. We investigated expression of cell entry machinery genes *ACE2, TMPRSS2, CTSB*, and *CTSL* by scRNAseq for different cell types of the human endometrium throughout the menstrual cycle^13^. scRNAseq was applied to endometrial samples from 27 healthy reproductive-age subjects in their natural menstrual cycles. For 19 participants, scRNAseq data was collected using the Fluidigm C1 system, resulting in 2,148 cells across the menstrual cycle. For the remaining 10 participants, scRNAseq data collection was performed using the 10x Chromium system, and analysis was done on a total of 71,032 cells targeting the preconception period of the menstrual cycle, when embryo implantation and decidualization occur^13^ (Supplementary Figure 1). For two women, we collected both C1 and 10x data, one from mid-secretory phase and the other early-secretory (**Supplementary Fig. 1B**).

scRNAseq analysis across the menstrual cycle (n=2,148 cells) showed low expression of *ACE2* in all cell types across the cycle (up to ~0.3 % of stromal fibroblasts or ~1 % of unciliated epithelia) (Figure 1). *TMPRSS2* was expressed mostly in glandular epithelial cells (12%) during early proliferative phase. Interestingly, *CTSB* and *CTSL* were expressed in both stromal and epithelial cells across all phases of the menstrual cycle, with *CTSB* more abundant than *CTSL* and *CTSL* more abundantly expressed in stromal fibroblasts than in unciliated epithelia. Co-expression of *ACE2* and *TMPRSS2*, *ACE2* and *CTSB*, or *ACE2* and *CTSL* was residual (0.09%; 0.23% and 0.14 % respectably) in all phases of the menstrual cycle (Supplementary Figure 2).

**Figure 1:**
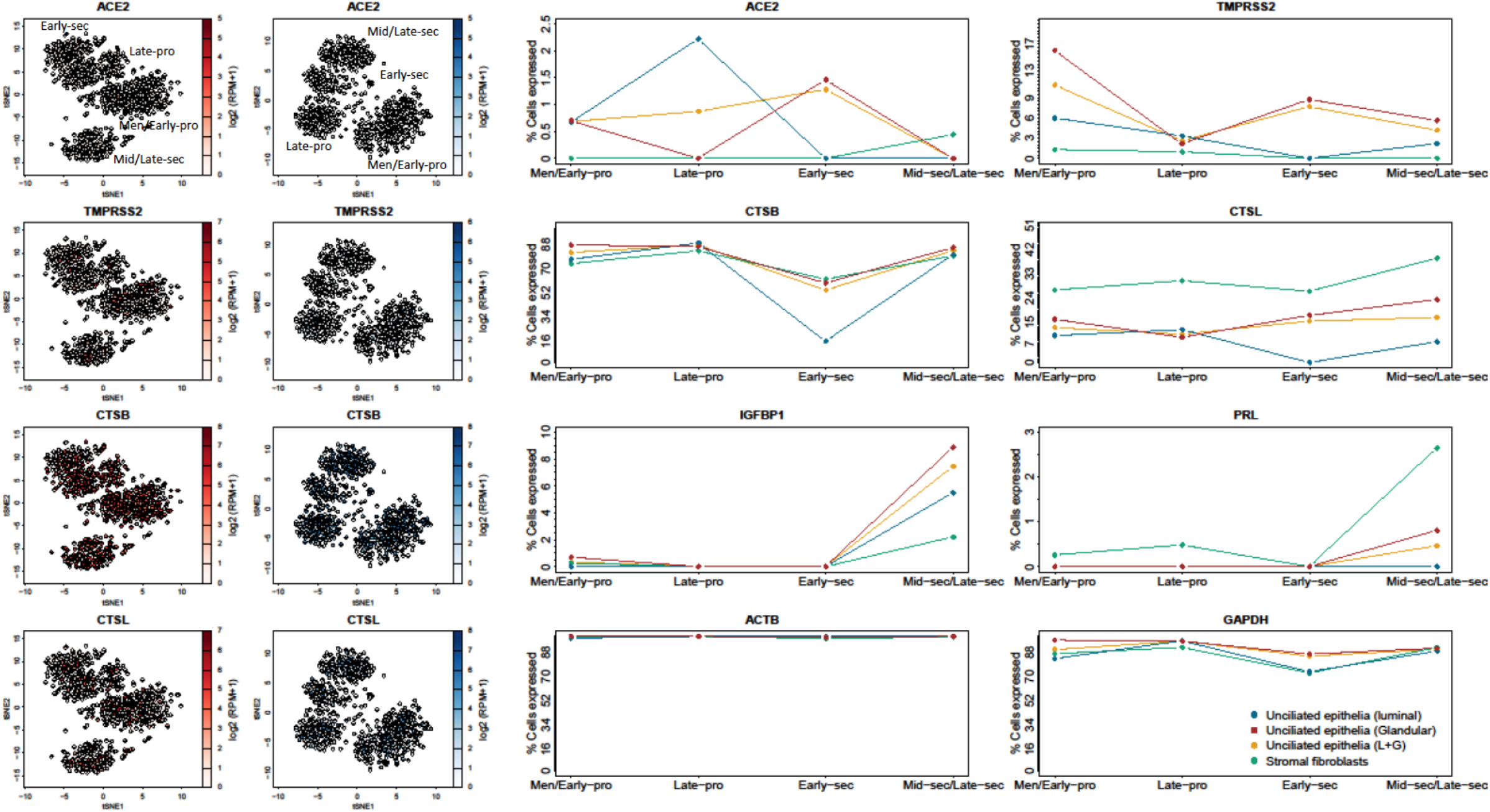
Dynamics of abundance of cells expressing *ACE2, TMPRSS2, CTSB* and *CTSL* across the menstrual cycle in uncilated epithelia and stromal fibroblasts. Quantification was performed on the C1 dataset. IGFBP1 and PRL were used as decidualization markers of the stromal fraction. ACTB and GAPDH were used as housekeeping controls.

10x dataset (n=71,032 cells), with more donors sampled during the secretory phase of the cycle, recapitulated the results in the C1 dataset. During all secretory phases, *ACE2* expression was low and detected in <2% of unciliated epithelia and stromal fibroblasts (Figure 2). *TMPRSS2* was expressed in ~12% of unciliated epithelia in the proliferative phase and this abundance changed notably across the cycle, while this gene was consistenly lowly expressed in stromal fibroblasts. In contrast, *CTSB* was highly expressed in both epithelial and stromal cells (~50-90%). Same as in the C1 dataset, *CTSL* was more highly expressed in stromal fibroblasts (60%). In the preconception phase, co-expression of *ACE2* with *TMPRSS2* was observed only in unciliated epithelia, representing ~0.016% of this cell type, while co-expression of *ACE2* with *CTSB* and *ACE2* with *CTSL* occurred in ~ 0.25% and 0.13% of cells, respectively (Supplementary Figure 2).

**Figure 2:**
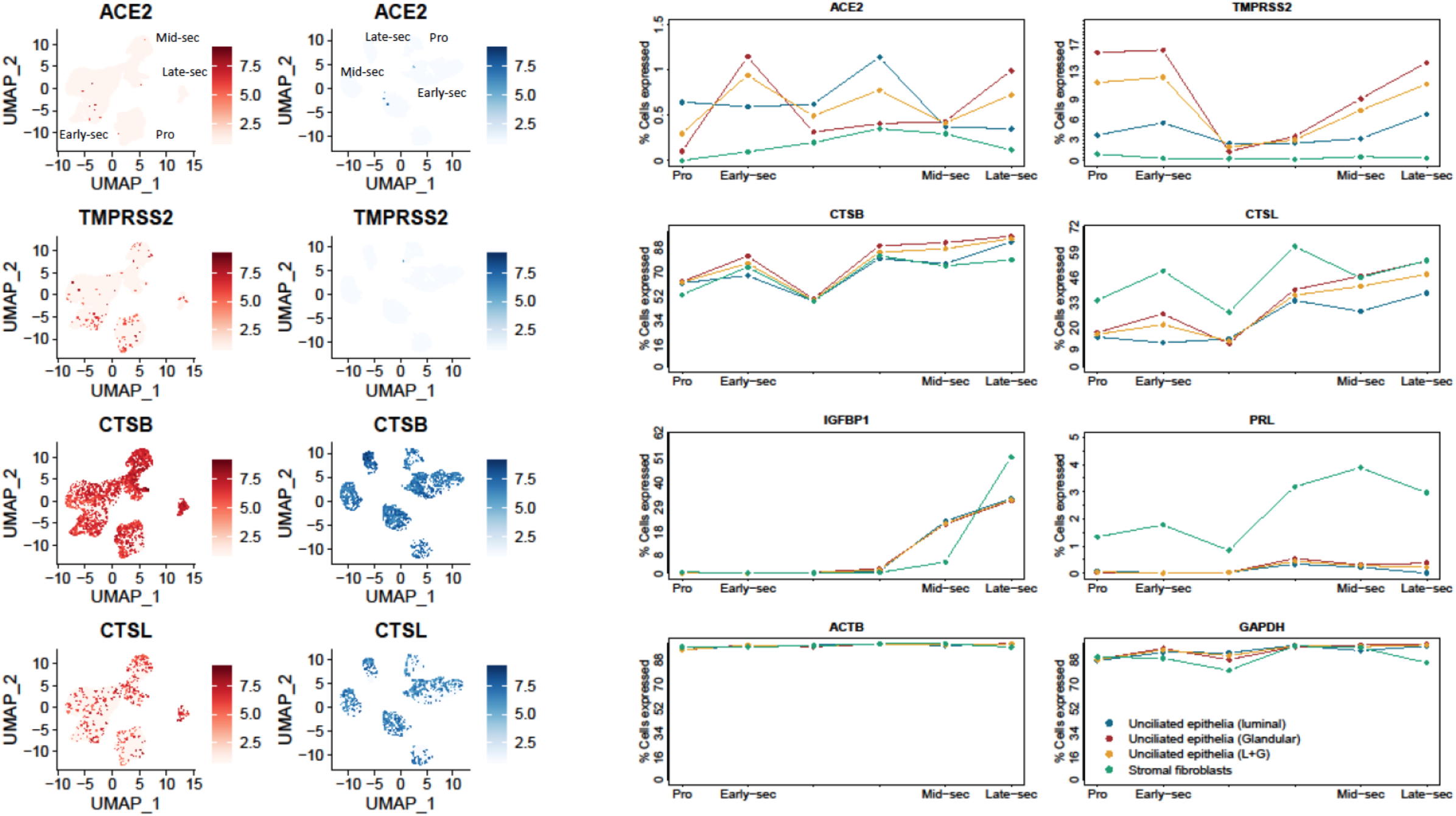
Dynamics of abundance of cells expressing *ACE2*, *TMPRSS2*, *CTSB* and *CTSL* across the menstrual cycle in uncilated epithelia and stromal fibroblasts. Quantification was performed on the 10X dataset. IGFBP1 and PRL were used as decidualization markers of the stromal fraction. ACTB and GAPDH were used as housekeeping controls.

## Discussion

In contrast to the reported expression of ACE2 in decidua and trophoblast cells in early pregnancy^14^, we did not detect higher *ACE2* transcripts in endometrial epithelial or stromal cells during the preconception period. High expression of *CTSB* and *CTSL* may imply that some part of the SARS-CoV-2 cell entry machinery is active in the endometrium at this time. Yet, because <1% of glandular epithelial cells simultaneously expressed all four genes of the cell entry machinery, our findings suggest low efficiency of SARS-CoV-2 viral infection in the human endometrium before conception. This study offers a useful resource to guide reproductive decisions when assessing the risk of endometrial infection by SARS-CoV-2 during the preconception period in asymptomatic COVID-19 carriers.

## Methods

### Subject details

All procedures involving human endometrium were conducted in accordance with the Institutional Review Board (IRB) guidelines for Stanford University under the IRB code IRB-35448 and IVI Valencia, Spain under the IRB code 1603-IGX-016-CS. Collection of endometrial biopsies was approved by the IRB code 1603-IGX-016-CS. There were no medical reasons to obtain the endometrial biopsies. Healthy ovum donors were recruited in the context of the research project approved by the IRB. Informed written consent was obtained from each donor in her natural menstrual cycle (no hormone stimulation) before an endometrial biopsy was performed. De-identified human endometrium was obtained from women aged 18-34, with regular menstrual cycle (3-4 days every 28-30 days), BMI ranging 19-29 kg/m^2^ (inclusive), negative serological tests for HIV, HBV, HCV, RPR and normal karyotype. Women with the following conditions were excluded from tissue collection: recent contraception (IUD in past 3 months; hormonal contraceptives in past 2 months), uterine pathology (endometriosis, leiomyoma, or adenomyosis; bacterial, fungal, or viral infection), and polycystic ovary syndrome.

### Endometrium tissue dissociation and population enrichment

Endometrial tissue digestion and scRNAseq library generation were previously described in 13.

A two-stage dissociation protocol was used to dissociate endometrium tissue and separate it into stromal fibroblast and epithelium enriched single cell suspensions. Prior to the dissociation, the tissue was rinsed with DMEM (Sigma) on a petri dish to remove blood and mucus. Excess DMEM was removed after the rinsing. The tissue was then minced into pieces as small as possible and dissociated in collagenase A1 (Sigma) overnight at 4 °C in a 50 mL falcon tube at horizontal position. This primary enzymatic step dissociates stromal fibroblasts into single cells while leaving epithelial glands and lumen mostly undigested. The resulting tissue suspension was then briefly homogenized and left un-agitated for 10 mins in a 50 mL Falcon tube at vertical position, during which epithelial glands and lumen sedimented as a pellet and stromal fibroblasts stayed suspended in the supernatant. The supernatant was therefore collected as the stromal fibroblast-enriched suspension. The pellet was washed twice in 50 mL DMEM to further remove residual stromal fibroblasts. The washed pellet was then dissociated in 400 μL TrypLE Select (Life technology) for 20 mins at 37°C, during which homogenization was performed via intermittent pipetting. DNaseI (100 μL) was then added to the solution to digest extracellular genomic DNA. The digestion was quenched with 1.5 mL DMEM after 5 min incubation. The resulting cell suspension was then pipetted, filtered through a 50 μm cell strainer, and centrifuged at 1000 rpm for 5 min. The pellet was re-suspended as the epithelium-enriched suspension.

### Fluidigm C1 single cell capture, imaging, and cDNA generation

For cell suspension of both portions, live cells were enriched via MACS dead cell removal kit (Miltenyi Biotec) following the manufacture’s protocol. The resulting cell suspension was diluted in DMEM into a final concentration of 300-400 cells/μL before being loaded onto a medium C1 chip for mRNA Seq (Fluidigm). Live dead cell stain (Life Technology) was added directly into the cell suspension. Single cell capture, mRNA reverse-transcription, and cDNA amplification were performed on the Fluidigm C1 system using default scripts for mRNA Seq. All capture site images were recorded using an in-house built microscopic system at 20x magnification through phase, GFP, and Y3 channels. 1μL pre-diluted ERCC (Ambion) was added into the lysis mix, resulting in a final dilution factor of 1:80,000 in the mix.

### Fluidigm C1 single cell RNAseq library generation

Single-cell cDNA concentration and size distribution were analyzed on a capillary electrophoresis-based automated fragment analyzer (Advanced Analytical). Tagmented and barcoded cDNA libraries were prepared only for cells imaged as singlet or empty at the capture site and with > 0.06 ng/uL cDNA generated. Library preparation was performed using Nextera XT DNA Sample Preparation kit (Illumina) on a Mosquito HTS liquid handler (TTP Labtech) following Fluidigm’s single cell library preparation protocol with a 4x scale-down of all reagents. Dual-indexed single-cell libraries were pooled and sequenced in pair-end reads on Nextseq (Illumina) to a depth of 1-2 ×10^6^ reads per cell. bcl2fastq v2.17.1.14 was used to separate out the data for each single cell by using unique barcode combinations from the Nextera XT preparation and to generate *.fastq files.

### Chromium 10x single cell capture and cDNA generation

Biopsies in the 10x dataset were obtained and dissociated following the same protocol used for the C1 dataset. They served as anchors for a direct comparison between the two datasets, and will be referred as “anchor biopsies” in this study. Following live cell enrichment via MACS (see description for the Fluidigm C1 dataset), live cells were washed twice with PBS to remove ambient RNA. The resulting epithelial and stromal portions were combined with a 1:1 ratio in concentration and were loaded onto the Chromium Next GEM Chip G (10x Genomics) for each donor. GEM generation & barcoding, reverse-transcription, cDNA generation, and library construction were done following the manufacture’s protocol (Single cell 3’ reagent kits v3.1, 10x Genomics). Dual-indexed single-cell libraries were pooled and sequenced in pair-end reads on Novaseq (Illumina).

### Quantification and statistical analysis

Single cell RNAseq data pre-processing, quality control and downstream statistical analysis were previously described in 13. Definition of endometrial cell types, states and temporal phases at single cell resolution followed that in 13. For quantifying percentage of cells expressing a gene in a cell type and/or an endometrial phase, a gene is considered expressed in a cell if its normalized count is higher than 0. All quantification was done in R.

## Acknowledgements

This study was jointly supported by the March of Dimes, Chan Zuckerberg Biohub and MINECO/FEDER (SAF-2015-67164-R, to C.S.) (Spanish Government). W.W. was supported by the Stanford Bio-X Graduate Bowes Fellowship and Chan Zuckerberg Biohub. F.V. was supported by the Miguel Servet Program Type II of ISCIII (CPII18/00020) and the FIS project (PI18/00957).

## Author contributions

F.V. contributed to conception, design, acquisition and interpretation of data and drafted the work. W. W. contributed to the acquisition, analysis, interpretation of data and drafted the work. I.M. contributed to design, and interpretation of data. S.R.Q. and C.S. contributed to conception, design, interpretation of data and drafted the work. All the authors have substantively revised the manuscript and approved the submitted version.

