## Supplementary Information for "Single-cell RNA Expression of SARS-CoV-2 Cell Entry Factors in Human Endometrium during Preconception"

Supplementary Figure 1: (A) Experimental design that represents the two methods used for single cell isolation and number of cells isolated. (B) Distribution of samples across the menstrual cycle day (number of days after the onset of last menstrual bleeding) and endometrial phases (assigned based on scRNAseq data) for the C1 (bottom) and 10x (top) dataset. Dots with * and # to the right are donors from whom both C1 and 10x data was collected. Phase 1-4: major endometrial phases identified in both unciliated epithelia and stromal fibroblasts using whole transcriptomic scRNAseq data.

Supplementary Figure 2: Percentage of cells that co-express *ACE2* and *TMPRSS2*; *ACE2* and *CTSB*; *ACE2* and *CTSL* across the menstrual cycle in the C1 dataset.

Supplementary Figure 3: Percentage of cells that co-express *ACE2* and *TMPRSS2*; *ACE2* and *CTSB*; *ACE2* and *CTSL* across the menstrual cycle in the 10X system.
