## Supplementary figures and images for "Single-cell RNA Expression of SARS-CoV-2 Cell Entry Factors in Human Endometrium during Preconception"

### Supplementary Figure 1

**Supplementary Figure 1**

**(A)**

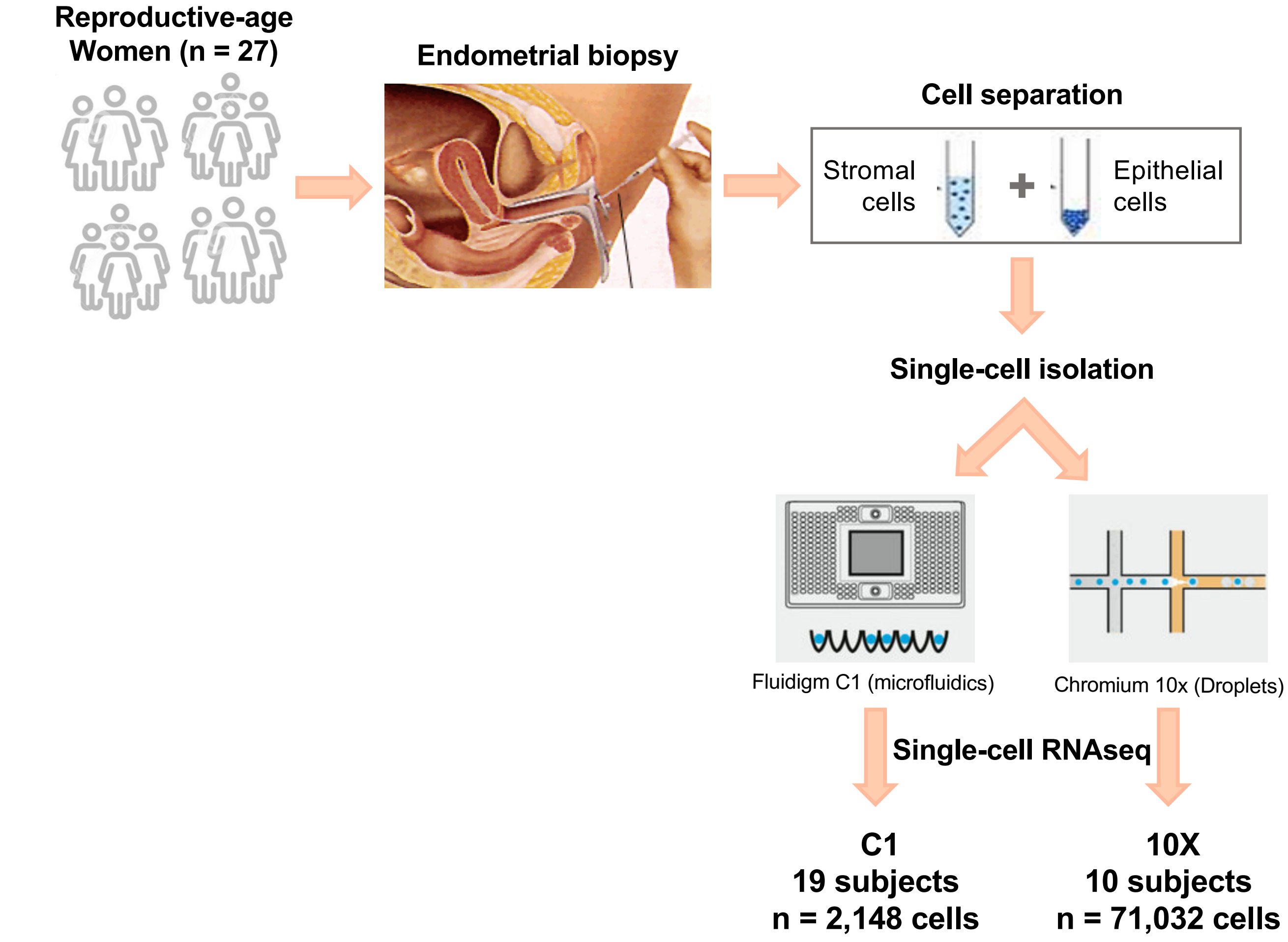


**(B)**

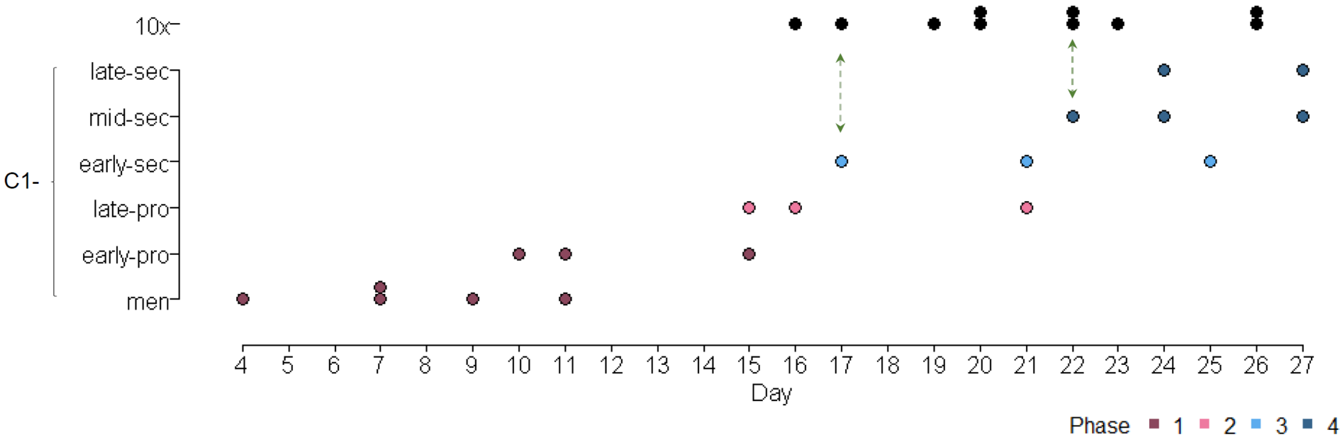
