## Supplementary Figure 2 for "Single-cell RNA Expression of SARS-CoV-2 Cell Entry Factors in Human Endometrium during Preconception"

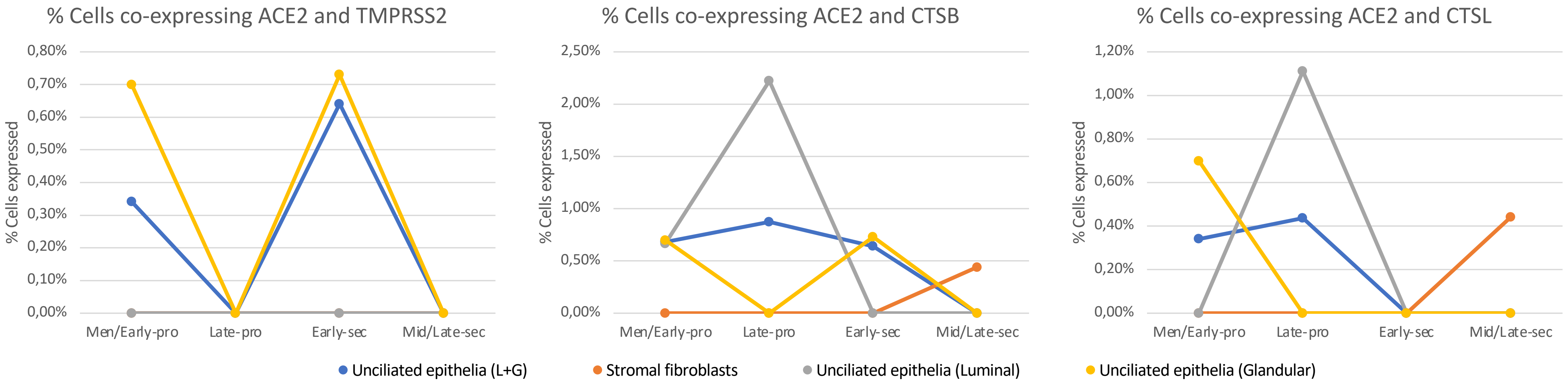


Total cell counts

| TMPRSS2 | Men/Early-pro | Late-pro | Early-sec | Mid/Late-sec |
|---|---|---|---|---|
| epi | 1 | 0 | 1 | 0 |
| str | 0 | 0 | 0 | 0 |
| epi_l | 0 | 0 | 0 | 0 |
| epi_g | 1 | 0 | 1 | 0 |

Total cell counts

| CTSB | Men/Early-pro | Late-pro | Early-sec | Mid/Late-sec |
|---|---|---|---|---|
| epi | 2 | 2 | 1 | 0 |
| str | 0 | 0 | 0 | 1 |
| epi_l | 1 | 2 | 0 | 0 |
| epi_g | 1 | 0 | 1 | 0 |

Total cell counts

| CTSL | Men/Early-pro | Late-pro | Early-sec | Mid/Late-sec |
|---|---|---|---|---|
| epi | 1 | 1 | 0 | 0 |
| str | 0 | 0 | 0 | 1 |
| epi_l | 0 | 1 | 0 | 0 |
| epi_g | 1 | 0 | 0 | 0 |

**Supplemental Figure 2**
