## Supplementary Figure 3 for "Single-cell RNA Expression of SARS-CoV-2 Cell Entry Factors in Human Endometrium during Preconception"

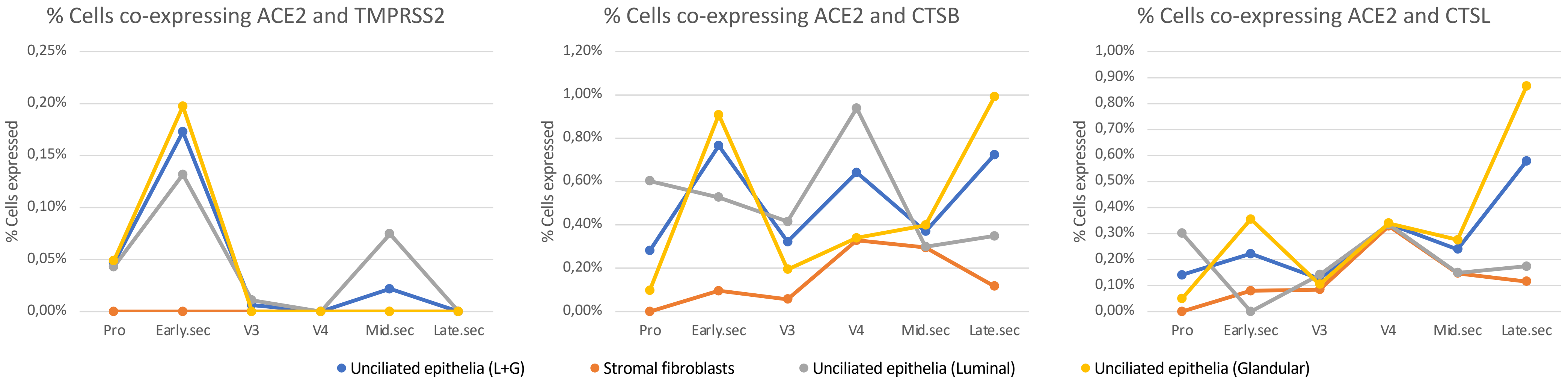


Total cell counts

| TMPRSS2 | Pro | Early.sec | V3 | V4 | Mid.sec | Late.sec |
|---|---|---|---|---|---|---|
| epi | 3 | 7 | 1 | 0 | 1 | 0 |
| str | 0 | 0 | 0 | 0 | 0 | 0 |
| epi_l | 1 | 2 | 1 | 0 | 1 | 0 |
| epi_g | 2 | 5 | 0 | 0 | 0 | 0 |

Total cell counts

| CTSB | Pro | Early.sec | V3 | V4 | Mid.sec | Late.sec |
|---|---|---|---|---|---|---|
| epi | 18 | 31 | 51 | 19 | 17 | 10 |
| str | 0 | 6 | 2 | 15 | 12 | 2 |
| epi_l | 14 | 8 | 38 | 14 | 4 | 2 |
| epi_g | 4 | 23 | 13 | 5 | 13 | 8 |

Total cell counts

| CTSL | Pro | Early.sec | V3 | V4 | Mid.sec | Late.sec |
|---|---|---|---|---|---|---|
| epi | 9 | 9 | 20 | 10 | 11 | 8 |
| str | 0 | 5 | 3 | 15 | 6 | 2 |
| epi_l | 7 | 0 | 13 | 5 | 2 | 1 |
| epi_g | 2 | 9 | 7 | 5 | 9 | 7 |

**Supplemental Figure 3**
